# Production of four-gene (GTKO/hCD55/hTBM/hCD39)-edited donor pigs and kidney xenotransplantation

**DOI:** 10.1101/2024.05.19.594896

**Authors:** Chang Yang, Yunfang Wei, Xinglong Li, Kaixiang Xu, Xiaoying Huo, Gang Chen, Heng Zhao, Jiaoxiang Wang, Taiyun Wei, Yubo Qing, Jianxiong Guo, Hongfang Zhao, Xiong Zhang, Deling Jiao, Zhe Xiong, Muhammad Ameen Jamal, Hong-Ye Zhao, Hong-Jiang Wei

## Abstract

**Background:** The number of multigene-modified donor pigs for xenotransplantation is increasing with the advent of gene editing technologies. However, which gene combination is suitable for which organ transplantation remains unclear.

**Methods:** In this study, we utilized CRISPR/Cas9 gene editing technology, PiggyBac transposon system and somatic cell cloning to construct GTKO/hCD55/hTBM/hCD39 four-gene-edited cloned (GEC) pigs and performed kidney transplantation from pig to rhesus monkey to evaluate the effectiveness of these GEC pigs.

**Results:** First, 107 cell colonies were obtained through drug selection, of which 7 were 4-GE colonies. Two colonies were selected for somatic cell nuclear transfer, resulting in 7 fetuses, of which 4 were GGTA1 biallelic knockout. Both fetuses had higher expression of hCD55, hTBM and hCD39. Therefore, these two fetuses were selected for two consecutive rounds of cloning, resulting in a total of 97 live piglets. After phenotype identification, the GGTA1 gene of these pigs was inactivated, and hCD55, hTBM and hCD39 were expressed in cells and multiple tissues. Furthermore, the numbers of monkey IgM and IgG binding to the peripheral blood mononuclear cells (PBMCs) of the 4-GEC pigs were markedly reduced. Moreover, 4-GEC porcine PBMCs had greater survival rates than those from wild-type pigs through complement-mediated cytolysis assays. In pig-to-monkey kidney xenotransplantation, the kidney xenograft successfully survived for 11 days. All physiological and biochemical indicators were normal, and no hyperacute rejection or coagulation abnormalities were found after transplantation.

**Conclusion:** These results indicate that the GTKO/hCD55/hTBM/hCD39 four-gene modification effectively alleviates immune rejection, and the pig kidney can functionally support the recipient monkey’s life.

## Background

Xenotransplantation is an effective way to solve the global organ shortage. With the rapid development of gene editing technology and new immunosuppressants, the United States of America (USA) has taken a leading step in conducting subclinical research on gene-edited (GE) pig-to-human organ transplantation. In 2022, the world’s first GE pig heart was transplanted into a human body at the University of Maryland Medical Center in the United States [1], which is a pioneer in the field of clinical xenotransplantation. In 2023, a kidney graft from a GE pig to a brain-dead patient was transplanted at New York University Langone Medical Center in the USA. This kidney functioned normally for 61 days (https://abcnews.go.com/Health/wireStory/pig-kidney-works-record-2-months-donated-body-103187024), thus establishing a new record for functional GE pig kidneys in humans.

Pig-to-human organ transplantation first requires genetic engineering of pigs to overcome immune rejection and molecular functional incompatibility. The α-galactose epitope synthesized by α1,3-galactosidyl transferase (GGTA1) induces hyperacute rejection, which is a major obstacle to xenotransplantation. Transplantation of pig organs with single-GE (GTKO) into nonhuman primates (NHPs) prolonged the survival of kidneys up to 68 days [2] and that of the heart up to 24 days [3]. Two recent cases of kidney transplantation from pigs to brain-dead humans also used GTKO pig kidneys, and these cases also confirmed the necessity of GGTA1 knockout [4]. Regardless of GGTA1, there are more than 30 other factors that have been related to xenograft rejection [5]. However, the current research still has demonstrated difficulty in defining the xenotransplantation outcomes between different genetically modified donor pigs, and which genetic combination is most suitable for which organ remains undetermined. Although David K.C. Cooper, a pioneering xenotransplantation researcher, believes that 9 GE donor pigs are sufficient to support xenotransplantation [6], some experts still recommend more careful consideration of the pros and cons of each edited gene with the objection that more is not always better.

In the case of GTKO, porcine xenografts are often attacked by non-Gal antibodies, which also leads to the activation of the complement system. CD55 can effectively inhibit the classic pathway and alternative pathway of the complement activation process by changing the activity of C3 and C5 convertases [7]. The survival time of GTKO/hCD55 GE pig kidney grafts transplanted into NHPs was prolonged to 499 days [8]. This greatly encouraged the use of other genetic modifications based on GTKO/hCD55, which could be a further breakthrough in the survival time of kidney xenografts from pigs to NHPs. The long-term survival of xenografts is often accompanied by the development of thrombotic microangiopathy, consumptive coagulopathy, or persistent inflammation that is mainly due to the incompatibility of coagulation molecules between pigs and humans [9]. The porcine thrombomodulin (TBM) complex cannot effectively activate human protein C, leading to graft thrombosis (Roussel et al., 2008). The overexpression of human TBM significantly increases activated protein C [10] and hence effectively inhibits thrombosis. In addition, TBM also protects vascular endothelial cells and inhibits neutrophil extracellular traps (NETs) and damage-associated molecular patterns (DAMPs) [11]. Moreover, CD39 can hydrolyze ATP, ADP, and AMP into adenosine, thereby exerting anti-inflammatory and antiplatelet aggregation effects [12]. Therefore, this study aimed to construct 4-GEC (GTKO/hCD55/hTBM/hCD39) xenotransplantation donor pigs and to conduct kidney transplantation research from pigs to NHP.

To ensure the long-term survival of porcine kidney xenografts in NHPs, first, cross-match between the donors and recipients is needed, and recipients with low antibody titers and low complement-dependent cytotoxicity for transplantation need to be selected [13]. Second, long-term survival depends on maintenance therapy with anti-CD40 mAb and anti-CD154 mAb-based immunosuppressive regimens [8, 14, 15]. Finally, long-term survival requires meticulous postoperative monitoring and care to ensure the maintenance of the recipient’s physiological indicators at normal levels and to take timely measures in life-threatening situations. This study used Cas9, PiggyBac transposon and somatic cell nuclear transfer (SCNT) technology to construct 4-GE (GTKO/hCD55/hTBM/hCD39) *Diannan* miniature pigs and performed kidney transplantation from pig to rhesus monkey to evaluate the effectiveness of these GE donor pigs.

## METHODS

Experimental animals used in this study and all surgical procedures were approved by the Animal Ethics Committee of Yunnan Agricultural University.

### Vector construction

To target porcine GGTA1 (GeneID: 396733) gene, two sgRNAs (sgRNA1: 5′-GCTACAGGCCTGGTGGTACA-3′; sgRNA2: 5′-GATGCGCATGAAGACCATCG-3′) were designed at the 8^th^ exon. After detecting the lack of single nucleotide polymorphisms (SNPs) in the target region, PX458-sgRNA vector was constructed. The hICAM2-hTBM-P2A-hCD39 expression framework driven by human ICAM2 promoter was constructed, and sub-cloned into the PiggyBac vector to obtain the hTBM-hCD39-piggyBac overexpression vector.

### Cell transfection, screening, and identification

The hCD55 overexpressed pig fetal fibroblasts (PFFs) transfected by using the hEF1α-hCD55-P2A-Puro piggyBac vector were thawed and cultured in DMEM containing 10% fetal bovine serum (FBS) for 1 day and cell growth status was observed. Cells, once attained logarithmic phase, were selected for electro-transfection. Approximately 7×10^5^ cells were suspended in the electro-transfection buffer containing PX458-sgRNA1 vector, PX458-sgRNA2 vector, hTBM-hCD39-piggyBac vector and piggyBac transposase vector at the ratio (1:1:1:2) and were incubated for 5 min, and electroporated by a Lonza 4D-nucleofector X Unit (AAF-1003X, Germany). After electroporation, cells were plated into T25 flasks for 24 h in DMEM supplemented with 10% FBS. Then, 3 μg/ml puromycin and 5 μg/ml hygromycin were added to the culture medium for 24−48 h to select successfully transfected cells. Subsequently, the survived cells were digested and ∼80 cells were seeded in a 100 mm diameter dish and cultured for 8 days to obtained single-cell colonies, which were further harvested for genotyping by PCR and Sanger sequencing using the primers shown in Table S1.

### Somatic cell nuclear transfer and embryo transfer

Oocyte collection, in vitro maturation, SCNT, and embryo transfer were performed as described previously [16]. Briefly, cultured cumulus-oocyte complexes (COCs) were isolated from cumulus cells by treating with 0.1% (w/v) hyaluronidase. The first polar body was enucleated via gentle aspiration using a beveled pipette in TLH-PVA, while the donor cells were injected into the perivitelline space of the enucleated oocytes. The reconstructed embryos were fused with a single direct current pulse of 200 V/mm for 20 µs using the Electro Cell Fusion Generator (LF201, NEPA GENE Co., Ltd., Japan) in fusion medium. Embryos were then cultured in PZM-3 for 0.5–1 h and activated with a single pulse of 150 V/mm for 100 ms in activation medium. The embryos were equilibrated in PZM-3 supplemented with 5 µg/ml cytochalasin B for 2 h at 38.5 °C in a humidified atmosphere with 5% CO_2_, 5% O_2_, and 90% N_2_ (APM-30D, ASTEC, Japan) and then cultured in PZM-3 medium with the same culture conditions described above until embryo transfer. The SCNT embryos were surgically transferred into the oviducts of the recipients, and a viable fetus was obtained through caesarian section after 31 days of pregnancy. A fetal fibroblast cell line was established, and subjected to PCR, T7EI enzyme digestion experiment, Sanger sequencing, flow cytometry and WB identification. Fetuses with biallelic knockout and target gene overexpression were selected for recloning. After completion of gestation period, GEC piglets were obtained through natural delivery and identified by the same way of fetus identification.

### Western blotting

Tissue samples were lysed in RIPA lysis buffer (Cat. no. BB⍰3201⍰2; Bestbio) with protease inhibitor (Cat. no. D1201⍰2; TransGen Biotech) at 4°C. Then, the total protein concentrations of the tissue lysates were determined using a BCA Protein assay kit (Cat. no. P0009⍰1 & 2; Beyotime institute of Biotechnology). Protein samples were then separated via 10% and 15% SDS–PAGE, and subsequently transferred to PVDF membrane. The membranes were incubated at room temperature (RT) for 30 min in 5% BSA solution with gentle shaking to block non⍰specific binding before incubation with the diluted primary antibody (hCD55, 1:3000, Cat. no. ab133684, Abcam; hTBM, 1:2000, Cat. no. sc-13164, Santa Cruz; hCD39, 1:3000, Cat. no. ab223842, Abcam). Subsequently, membranes were incubated with horseradish peroxidase⍰conjugated secondary antibody (hCD55 and hCD39, rabbit IgG, 1:5000 (Cat. no. HaF008, R&D Systems); hTBM and β-actin, mouse IgG, 1:10000 (Cat. no. HaF007, R&D Systems)) for 2 h at RT. Membranes were first washed three times and then incubated with reagent from an Easysee Western Blot kit (Cat. no. DW101-01, TransGen Biotech) and visualized using an imaging system (ChemiDoc^TM^ MP Imaging System, Bio⍰Rad Laboratories, Inc.).

### Quantitative polymerase chain reaction

Total RNA was extracted using TRIzol reagent (Cat. no. ET111-01-V2, TransGen Biotech) according to the manufacturer’s instructions, and the concentration and RNA quality were detected. Complementary DNA (cDNA) was synthesized from total RNA using a PrimeScript RT reagent Kit (Cat. no. RR047B, TaKaRa) and was used as a template to perform qPCR in TB green-based qPCR instrument (CFX-96, Bio-Rad, USA). The reaction was performed in a 20 µl reaction mixtures comprising 10 µl of 2×TB Green® Premix Ex Taq™ (Tli RNaseH Plus) (Cat. no. RR420A, TaKaRa), 1 µl of cDNA, 1 µl of forward primer, 1 µl of reverse primer, and 7 µl of ddH_2_O (primers listed in Table S1). The reaction program is as follows: 95°C for 30 s, followed by 40 cycles of 95°C for 10 s, and 62°C for 45 s. Three technical replicates were conducted for each sample and the relative expression levels of target genes were quantified by 2^-ΔΔct^.

### Immunofluorescence

The paraffin-embedded tissue blocks were cut into 5 µm, transferred to glass slides, dewaxed using xylene and gradient alcohol, put into a microwave oven to retrieve antigens with EDTA buffer (Cat. no. G1207, Servicebio Bio) at 92–98°C for 15 min, and cooled at room temperature. Then, sections were washed with PBS for three times (each time 3 min), incubated with autofluorescence quencher A (Cat. no. G1221, Servicebio Bio) at RT in dark for 15 min, washed with PBS for three times again, incubated with FBS at RT for 30 min, and dried. The dried sections were incubated with corresponding antibodies (αGal, Cat. no. ALX-650-001F-MC05, Enzo; hCD55, Cat. no. ab230638, Abcam; hTBM, Cat. no. sc-13164, Santa Cruz; hCD39, Cat. no. ab223842, Abcam). For visualization of hCD55, hTBM, hCD39 protein, Cy3-conjugated secondary antibodies (hCD55, Cat. no. ab230638, Abcam; hTBM, Cat. no. sc-13164, Santa Cruz; hCD39, Cat. no. ab223842, Abcam) were diluted with PBS containing 10% FBS (v/v = 1:200) and used to incubate sections at 4°C in dark for 2 h, and a negative control was incubated with PBS containing 10% FBS. Then, sections were washed with PBS three times and stained with DAPI (Cat. no. G1012, Servicebio Bio) for 3 min. After washing with PBS for 1 min, autofluorescence quencher B (Cat. no. G1221, Servicebio Bio) was added for 5 min and washed three times again. Finally, sections were mounted with anti-fluorescence quencher (Cat. no. G1401, Servicebio Bio) and imaged using an OLYMPUS BX53 fluorescence microscope.

### Cell flow cytometry

All flow cytometry assays were performed by CytoFLEX flow cytometer (Beckman Coulter, USA). For detection of GGTA1, hCD55, hTBM, and hCD39 surface molecules on fibroblasts derived four GE pigs and WT, cells were cultured in cell medium (Cat. no. TM002, ABM) containing 10% FBS (Cat. no. VS500T, Ausbian) and 1% Pen-Strep solution (Cat. no. 03-031-1BCS, Biological Industries) for 3 days. After collection, 1×10^5^ cells were incubated with mouse αGal (1:50, Cat. no. ALX-650-001F-MC05, ENZO SWIT), CD55 (1:5, Cat. no. 555694, BD USA), TBM (1:20, Cat. no. 564123, BD USA) and mouse CD39 (1:5, Cat. no. 560239, BD USA) for 1 h at 4℃, respectively. After that, cells were washed twice with PBS and centrifuged at 1400 rpm for 5 min. All data acquired from flow cytometry were analyzed by using FlowJo VX software.

### H&E staining

Tissue samples were fixed in 4% paraformaldehyde for 48–72 h, processed by an automatic tissue processor (Yd-12p, Jinhua Yidi, medical appliance Co., Ltd, China) and embedded in a paraffin block (Yd-6D, Jinhua Yidi, medical appliance Co., Ltd, China). The paraffin blocks were cut into 5-um-thick sections using a Microm HM 325 microtome (Thermo Scientific, USA) and allowed to dry on glass slides overnight at 37°C. Thereafter, the tissue sections were deparaffinized in xylene and rehydrated through graded ethanol dilutions. Sections were stained with hematoxylin–eosin (H&E) (Cat. no. G1120, Solarbio) according to manufacturer’s instruction and quantified.

### Rhesus monkey serum mediated-antibody binding assay

Rhesus monkey serum was collected and inactivated at 56℃ for 30 min, and was diluted 1:4 in staining buffer (PBS containing 1% FBS). And PFFs derived from four GE pig and WT pig were collected, washed twice and resuspended in staining buffer. 1×10^5^ cells were incubated with 100 µl inactive rhesus monkey serum (test group) or 100 µl PBS (negative control) for 45 min at RT. Then cells were washed with cold staining buffer to terminate reaction and incubated with goat anti-human IgM-FITC (1:100 dilution, Cat. no. A18842, Invitrogen), and IgG-FITC (1:200 dilution, Cat. no. 628411, Invitrogen) for 30 min at 4℃. Finally, cells were washed with cold staining buffer twice, centrifuged at 400×g for 4 min, and resuspended with 200 µl PBS for a CytoFLEX flow cytometer (Beckman Coulter, USA).

### Complement-dependent cytotoxicity assay

Rhesus monkey serum was collected and inactivated at 56℃ for 30 min, and was diluted 1:1 in staining buffer (PBS containing 1% FBS). And peripheral blood mononuclear cells (PBMCs) derived from four GEC pig and WT pig were collected, washed twice and resuspended in staining buffer. 1×10^5^ cells were incubated with 50 µl inactive rhesus monkey serum (test group) or 50 µl PBS (negative control) for 30 min at RT. Then cells were washed with cold staining buffer to terminate reaction and incubated with rabbit complement sera (1:3 dilution, Cat. no. S7764, Sigma) for 30 min at RT. Finally, cells were stained with PI staining solution (Cat. no. GA1174, Servicebio Bio) for 2 min, and analyzed for cell death using a CytoFLEX flow cytometer (Beckman Coulter, USA).

### Droplet digital PCR (ddPCR) for transgenic copy numbers

The samples including the human blood (positive control), the pre-transfected porcine cells (negative control), the selected cell lines (C21 and C78), the fetal cells (F02 and F06) and the kidney tissues of the cloned pig derived from fetuses F02 and F06 were collected and extracted DNA. Then, these DNA samples were digested by MseI restriction endonuclease (Cat. no. R0525, New England BioLabs) at 37 °C for 1 h, and inactivated at 65 °C for 10 min. The digested product was diluted to 5ng/ul and used as a ddPCR template. The primers and probes corresponding to hCD55, hCD39, hTBM, and GAPDH genes (Table S1), ddPCR super mix (Cat. no.1863024, Bio-Rad), and DNA template were mixed to prepare a 25ul reaction system. Then, ddPCR droplets were generated with QX100 Droplet Generator (Bio-Rad) and transferred to a 96 well plate. PCR reactions were performed at 95 °C for 10 min, 94 °C for 3 s, 56 °C for 1 min (39 cycles), and 98 °C for 10 min. After the reaction, the 96 well plate was placed in Bio-Rad Droplet Reader and QunataSoft Software was used to set up the experimental design and read the experiment. Once the program was finished, the copy numbers were analyzed through gating, according to the manufacturer’s instructions (Bio-Rad).

### Pig-to-nonhuman primate kidney transplantation

We performed pig-to-non human primate kidney transplantation at the Yunnan Province Xenotransplantation Research Engineering Center, Yunnan Agricultural University, Kunming, Yunnan, China. The immunosuppressive regimen and surgical procedures used are shown in Figure S1. First, pig PBMCs and rhesus monkey serum were used for cross-matching to select recipient monkeys with low antibody titers and low complement-dependent toxicity for kidney transplantation. Three days prior to surgical interventions, dexamethasone and promethazine were injected once a day to reduce the occurrence of allergic reactions caused by immunosuppressants. Antithymocyte globulin (ATG), anti-CD20 mAb, and anti-CD154 mAb were used to induce immunosuppression on days -3, -2, and -1, respectively, and the whole blood of the recipient monkey was collected for hematological analysis.

On the day of transplantation, the 4-GE donor pig underwent unilateral nephrectomy (left kidney); the recipient rhesus monkey received anti-CD154 mAb, tacrolimus, and methylprednisolone. Afterward, a left nephrectomy was performed, and a kidney from the donor pig was transplanted, followed by a right nephrectomy.

After surgical interventions, the recipient received an immunosuppressive regimen based on anti-CD154 mAb (5c8, NIH Nonhuman Primate Reagent Resource) supplemented by low molecular weight heparin, erythropoietin β, famotidine, aspirin, and ceftriaxone sodium or cefsulbactam sodium for adaptive therapy (Figure S1). Postoperative care provided by postgraduate students majoring in veterinary medicine included routine blood tests and blood creatinine, urine creatinine, albumin and other biochemical tests every two days to evaluate the degree of immune rejection, coagulation status, physiological status, renal and liver function, etc., of the recipient monkey.

### Statistical analysis

All data were analyzed using SPSS 22.0 software package (IBM Crop, Armonk, NY) and expressed as mean ± standard deviation of mean (SD). Statistical significance was defined as *P < 0.05, **P < 0.001.

## RESULTS

### Generation of the GTKO/hCD55/hTBM/hCD39 4-GEC pigs

We designed two sgRNAs that simultaneously targeted the GGTA1 gene in *Diannan* miniature pigs. Moreover, the expression of hTBM and hCD39 was driven by the human endothelial cell-specific promoter ICAM2, and a P2A element with a high self-cleavage rate was used to link hTBM and hCD39, constructing the corresponding PiggyBac transposon vector (Figure 1A). Cas9, sgRNA and hTBM/hCD39 overexpression vectors were cotransfected into PFFs overexpressing hCD55. After antibiotic selection, 107 cell colonies were obtained, and of these, 7 colonies (C17#, C21#, C26#, C29#, C57#, C63# and C78#) carried GGTA1 gene biallelic mutations (Figure 1B). Moreover, hTBM and hCD39 from these cell colonies were successfully integrated into the pig genome (Figure 1C). Based on the cell growth status, we selected C21# and C78# colonies as donor cells for the first round of cloning and transferred them into four surrogate sows. This resulted in a total of seven 31-day-old fetuses (Figure 2A). Among these seven fetuses, all carried hCD55, hTBM and hCD39 transgenes and GGTA1 mutations (Figure 2 B-D). The F01, F02, F03, and F06 fetuses exhibited biallelic mutations in GGTA1, as determined by Sanger sequencing (Figure 2E). However, hCD55, hTBM and hCD39 protein expression was detected in only the F02 and F06 fetuses (Figure 2F). Hence, the fibroblasts from these two fetuses were selected as donor cells for the second round of cloning, and 42 and 16 viable piglets were obtained, respectively (Figure 3A). Additionally, we selected 3 cloned piglets for the third round of cloning and obtained 39 live piglets through natural delivery.

**Figure 1.**
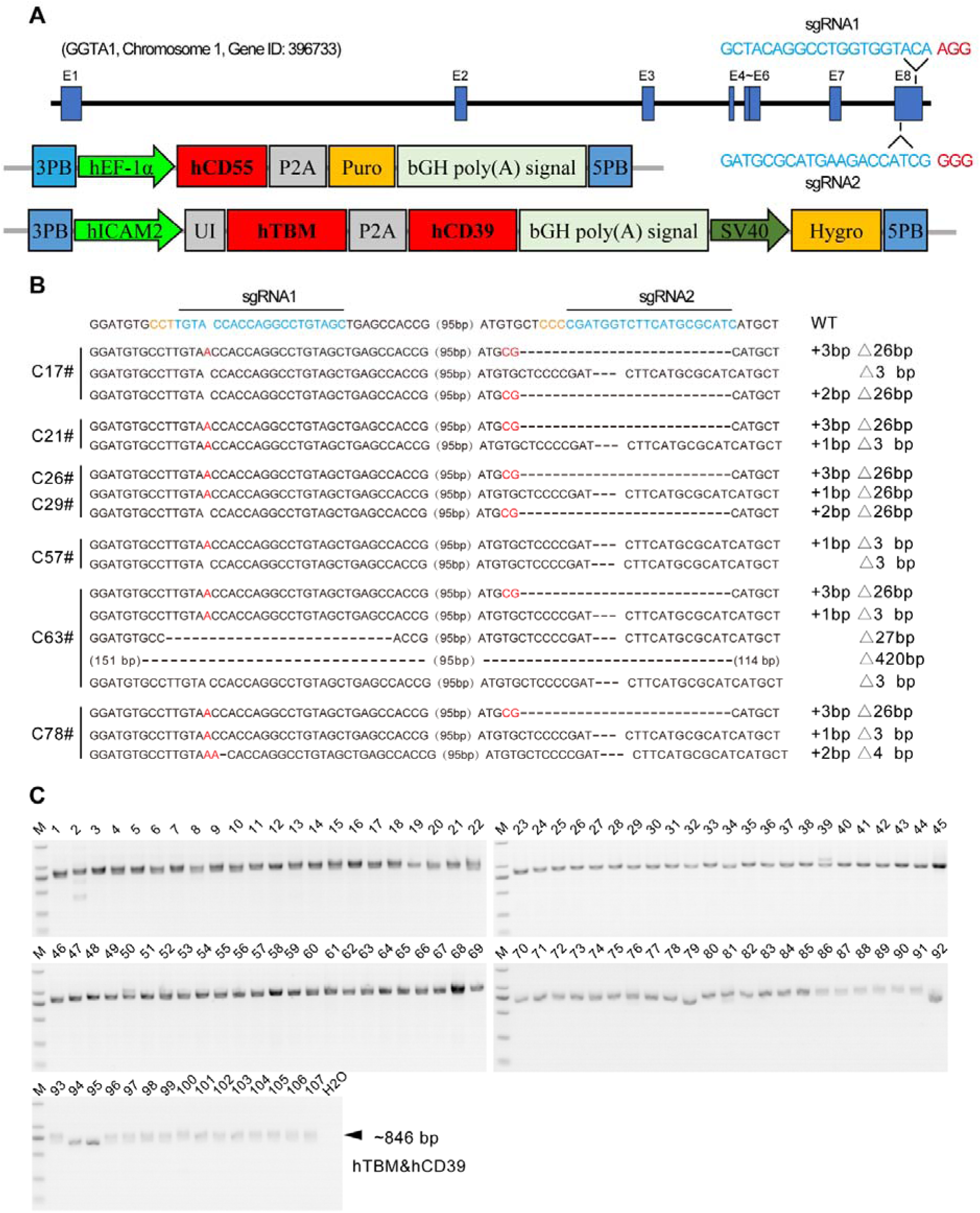
Construction of gene editing vectors for GGTA1 knockout, hCD55, hTBM and hCD39 overexpression as well as screening out of cell colonies A. Knockout of porcine GGTA1 gene by CRISPR/Cas9 targeting exon 8. The expression of cDNA of human CD55 gene was driven by human EF-1α promoter, and the expression of cDNAs of hTBM and hCD39 gene were driven by human ICAM2 promoter. B. The sequence of the targeting region of GGTA1 gene in cell colonies by Sanger sequencing after transfection and drug selection. C. hTBM and hCD39 genes were successfully intergrated into pig genome in cell colonies by PCR identification. All of 107 cell colonies were positive for transgene integration.

**Figure 2.**
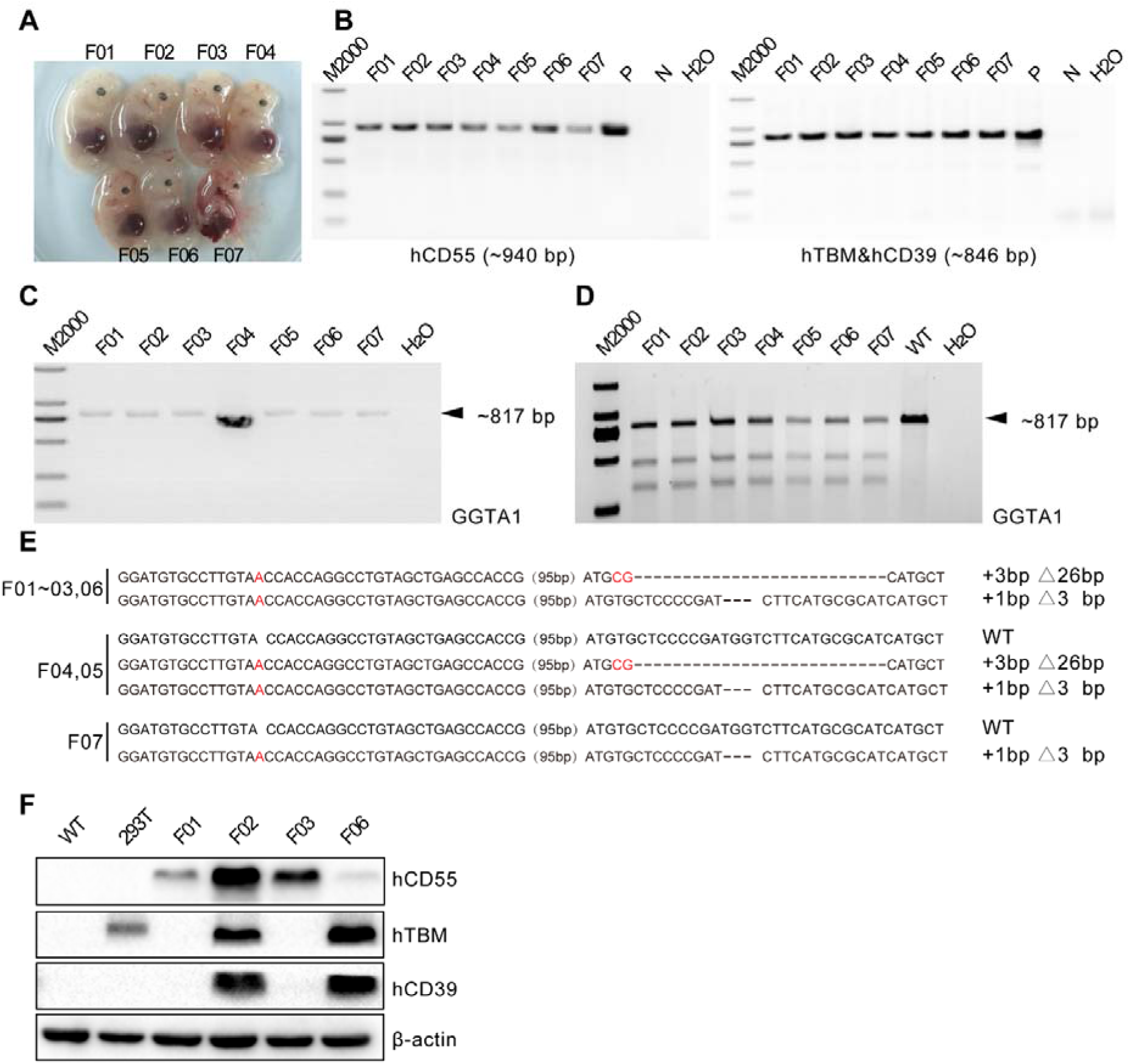
Generation and identification of gene etiting cloned fetuses. A. Photo of 7 cloned fetuses obtained after 33-day pregnancy. B. The integration of hCD55, hTBM and hCD39 genes into fetal genome was confirmed by PCR. C. The PCR products of targeting region of GGTA1 gene in fetuses. D. The mutation of GGTA1 gene in fetuses was confirmed by T7ENI cleavage assay. E. The sequence of the targeting region of GGTA1 gene in fetuses by Sanger sequencing. F. The protein expression of hCD55, hTBM and hCD39 genes in fetuses was confirmed by Western Blot.

**Figure 3.**
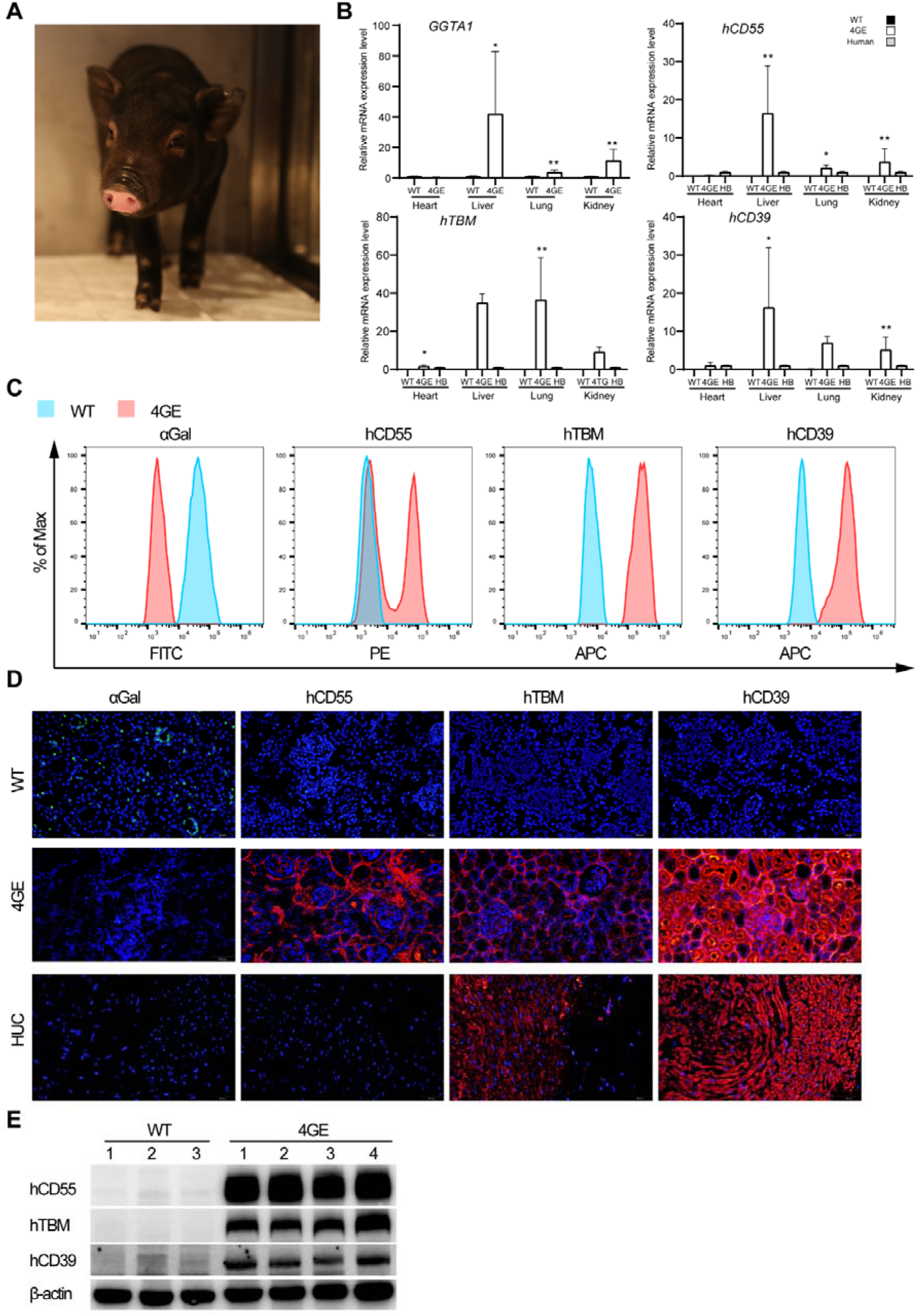
Phenotype of GTKO/hCD55/hTBM/hCD39 four-gene edited cloned pigs. A. Photo of GEC piglet. B. The mRNA expression levels of GGTA1, hCD55, hTBM and hCD39 in heart, liver, lung and kidney tissues of GEC pigs (n=3). WT, wild type; HB, human blood. C. The expression of αGal, hCD55, hTBM and hCD39 gene in GEC porcine fibroblasts was confirmed by flow cytometry. D. The protein expression of αGal, hCD55, hTBM and hCD39 gene in GEC porcine kidney tissues was confirmed by immunofluorescence. HUC, human umbilical cord. E. The protein expression of hCD55, hTBM and hCD39 gene in GEC porcine kidney tissues was confirmed by Western Blot.

### Phenotype identification of the GTKO/hCD55/hTBM/hCD39 4-GEC pigs

We first determined the genotype of the targeting region at the GGTA1 gene in cloned piglets, and the results showed that these piglets carried a biallelic mutation in GGTA1, consistent with the genotype of fetuses F02 and F06 (Table S2). Then, we measured the mRNA expression levels of the GGTA1, hCD55, hTBM and hCD39 genes in the heart, liver, lung and kidneys of cloned piglets. The mRNA expression levels of GGTA1 in the liver, lung and kidneys of the 4-GE pigs were significantly increased, while the expression levels of the hCD55, hTBM and hCD39 genes were higher than those of human whole blood (Figure 3B). Moreover, the αGal epitope was deleted in the 4-GEC pigs, and three protein molecules, hCD55, hTBM, and hCD39, were detected through flow cytometric analysis of fibroblasts (Figure 3C). Immunofluorescence and western blot analysis also confirmed that the αGal epitope was deficient, and hCD55, hTBM and hCD39 were expressed in the kidneys of the 4-GEC pigs (Figure 3D and E). In addition, we confirmed the copy numbers of hCD55, hTBM and hCD39 transgenes in donor cell lines, cloned fetuses and pig. The results showed that the copy numbers of hCD55 in donor cell line C21 and C78 were 7 and 5 respectively, while in fetuses and pigs, the copy numbers were 10 (Figure 4A). The copy numbers of hTBM in donor cell line C21 and C78 were 11 and 5 respectively, while in fetuses and pigs, the copy numbers varied from 12 to 15 (Figure 4B). The copy numbers of hCD39 in donor cell line C21 and C78 were 12 and 5 respectively, while in fetuses and pigs, the copy numbers varied from 12 to 15 (Figure 4C). Overall, the copy numbers of transgenes in fetuses and pigs were higher than those of donor cells.

**Figure 4.**
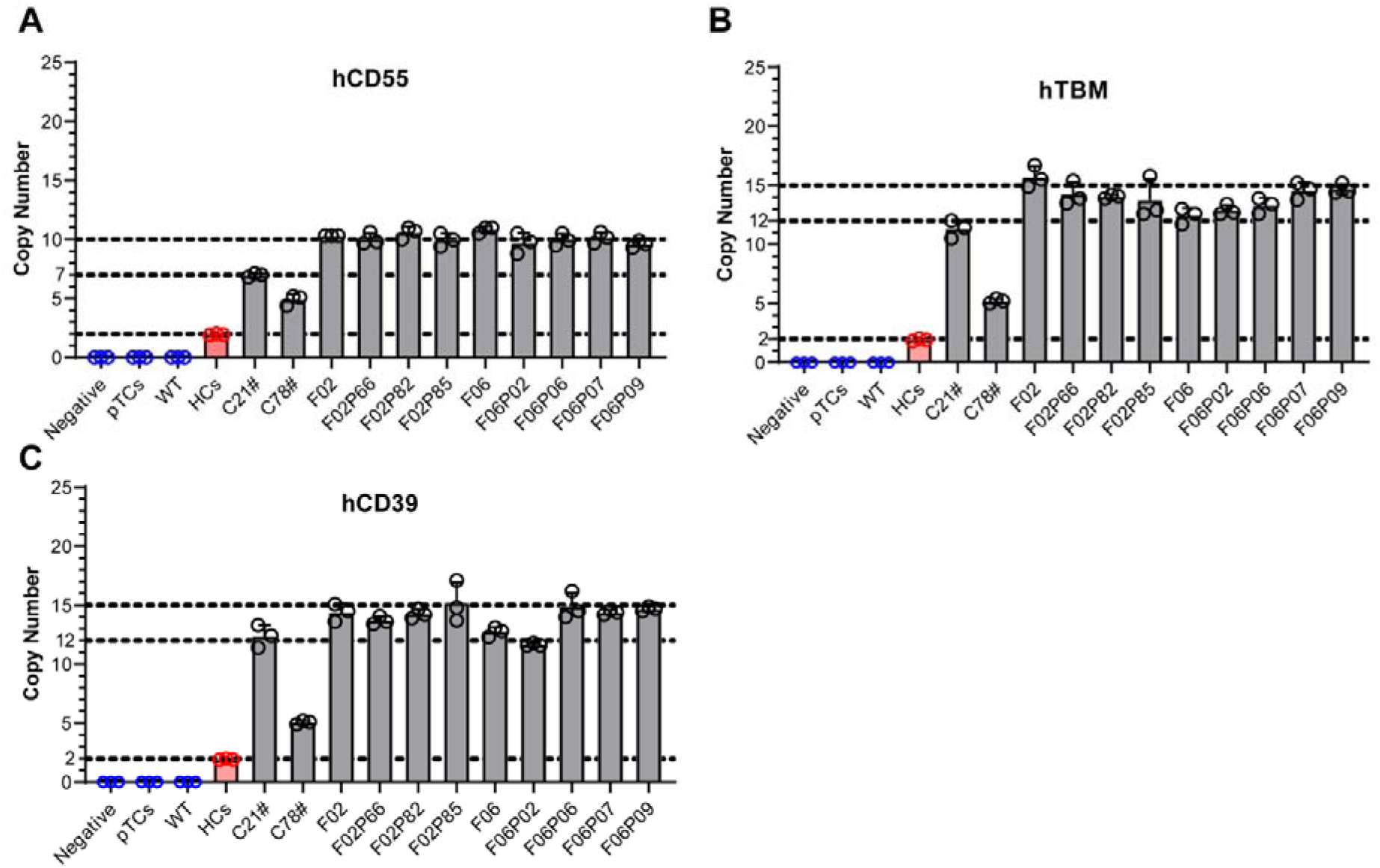
Copy numbers of transgenes hCD55, hTBM and hCD39 in donor cell lines, cloned fetuses and pigs. The copy numbers of transgenes hCD55 (A), hTBM (B) and hCD39 (C) were detected by ddPCR. Negative, water; pTCs, porcine pre-transfected cells; WT, wild-type pig kidney; HCs, human cells.

### Crossmatch between the GTKO/hCD55/hTBM/hCD39 4-GEC pigs and rhesus monkeys

Before transplanting kidneys from pigs to NHPs, we collected serum from 12 rhesus monkeys and conducted cross-matching experiments with 4-GEC pig PBMCs. The numbers of IgG and IgM binding to PBMCs from a 4-GEC pig (F02P85) to monkeys were significantly reduced compared to those of WT pigs (Figure 5A and B). Complement-dependent cytotoxicity experiments further confirmed that the four gene modifications protected porcine cells from complement attack, thereby improving the survival rate of the porcine cells (Figure 5C). Among the monkeys, monkey No. 20 had the lowest IgM titer (Figure 5D), but monkey No. 28 had the lowest IgG titer and anticomplement-dependent cytotoxicity (Figure 5E and F). Thus, No. 28 was selected as the recipient for kidney xenotransplantation.

**Figure 5.**
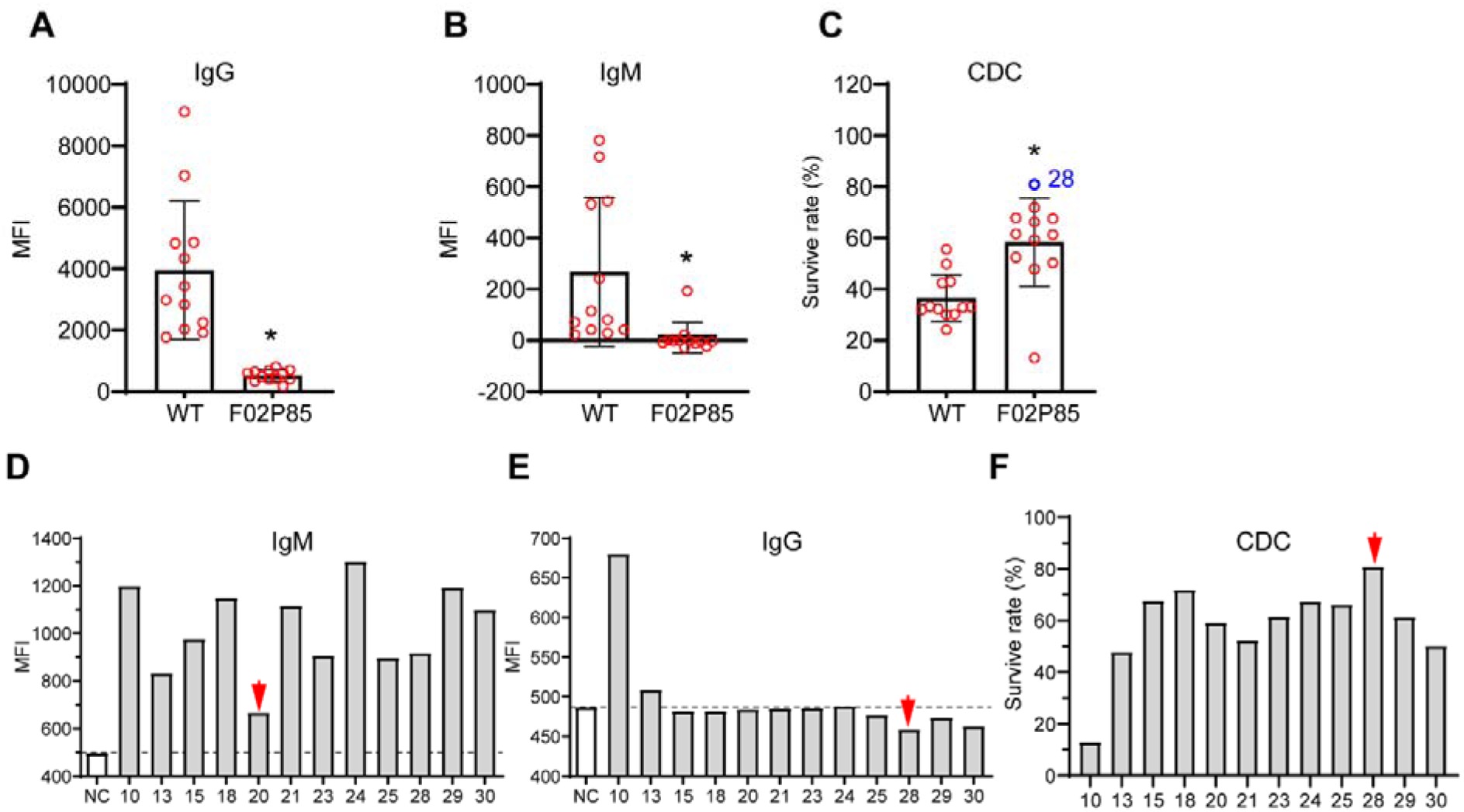
Crossmatch between the GTKO/hCD55/hTBM/hCD39 4-GEC pigs and rhesus monkeys. A. The levels of monkey IgG binding to 4-GEC porcine peripheral blood mononuclear cells (PBMCs). B. The levels of monkey IgM binding to 4-GEC porcine PBMCs. C. The survival rate of 4-GEC porcine PBMCs after 50% monkey serum incubation for 2 hours. The PBMCs of monkey 28# had the highest survival rate and was selected as recipient. D. The levels of each monkey IgG binding to 4-GEC porcine PBMC. E. The levels of each monkey IgM binding to 4-GEC porcine PBMC. F. The survival rate of 4-GEC porcine PBMCs after incubation with each monkey serum.

### Kidney xenotransplantation from the 4-GEC pig to a rhesus monkey

Before transplanting the kidney xenograft, the recipient monkey was immunosuppressed with ATG, anti-CD20, and anti-CD154 monoclonal antibodies. After transplantation, an immunosuppressive regimen based on methylprednisolone and tacrolimus combined with an anti-CD154 monoclonal antibody was used as maintenance treatment. The recipient monkey survived for 11 days, and no HAR was observed. After autopsy, it was observed that the kidney from the 4-GEC pig was congested, swollen, and necrotic (Figure 6A). H&E staining revealed bleeding in some parts of the pig kidney and extravasation of red blood cells, but the glomerular structure remained relatively intact (Figure 6B). In the postoperative period, there were no significant changes in the immunoglobulins IgA, IgG and IgM (Figure 6C-E) and complement C3 and C4 concentrations (Figure 6F and G) in the serum of the recipient monkey. Immunohistochemical analysis showed a small amount of complement C3c deposition in the pig kidney (Figure 6H).

**Figure 6.**
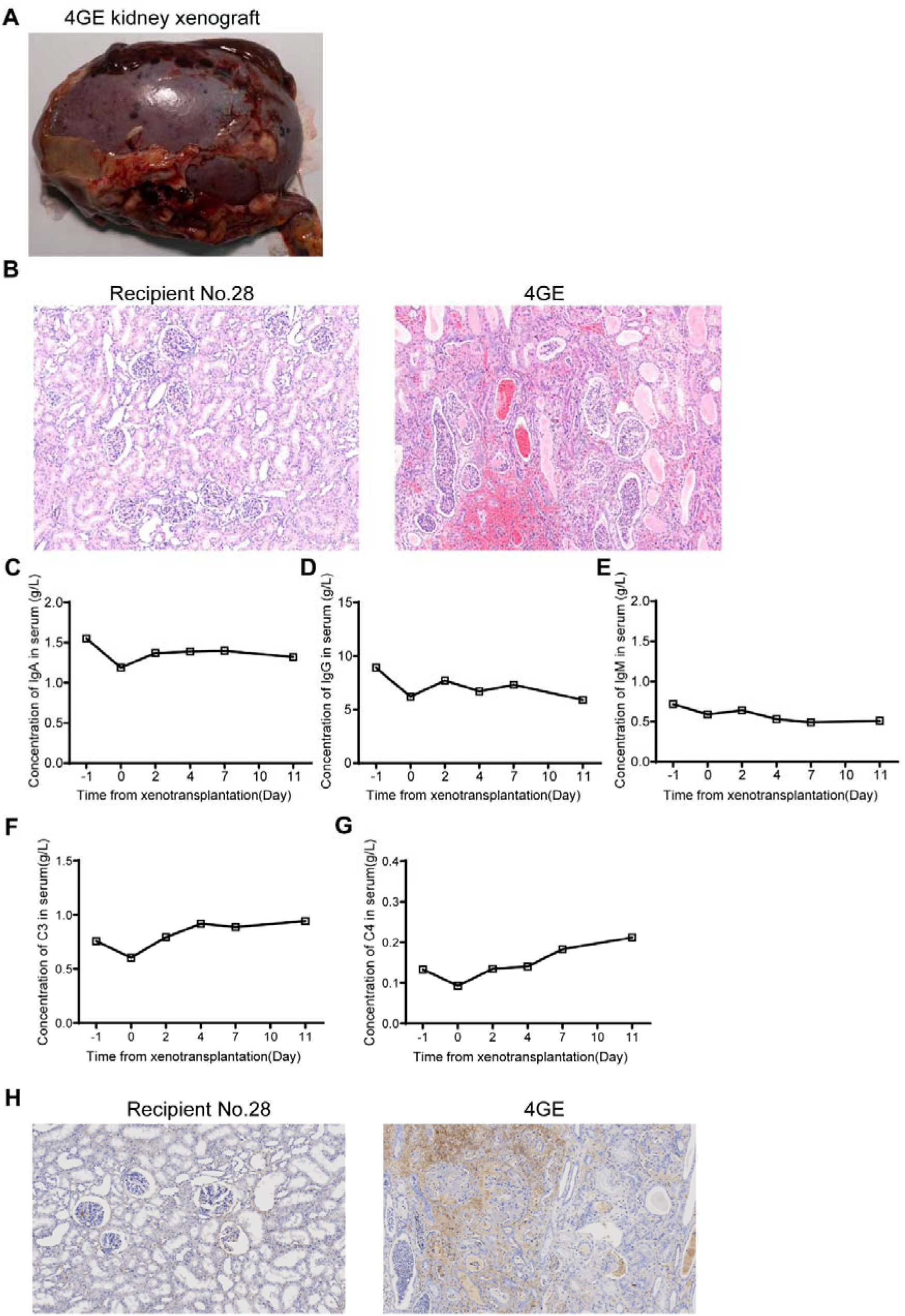
Pathological analysis of kidney xenograft and humoral immunity of recipient monkey. A. Photos of kidney excised from recipient monkey and porcine kidney after function loss. B. H&E staining of recipient monkey and porcine kidney xenograft. C-G. The levels of IgA (C), IgG (D), IgM (E), complement C3 (F) and C4 (G) in serum of recipient monkey. in serum of recipient monkey. H. Complement C3c deposition in porcine kidney xenograft confirmed by immunohistochemistry.

We monitored changes in the number of white blood cells, neutrophils, basophils, eosinophils, lymphocytes, and monocytes of recipient monkey. The results indicated that the number of leukocytes and neutrophils was abnormally elevated 7 days post-transplantation. The number of basophils and eosinophils was relatively stable, while the number of lymphocytes and macrophages tended to increase (Figure 7A-F). Specifically, macrophage infiltration was observed in the kidney xenograft after functional loss (Figure 7G).

**Figure 7.**
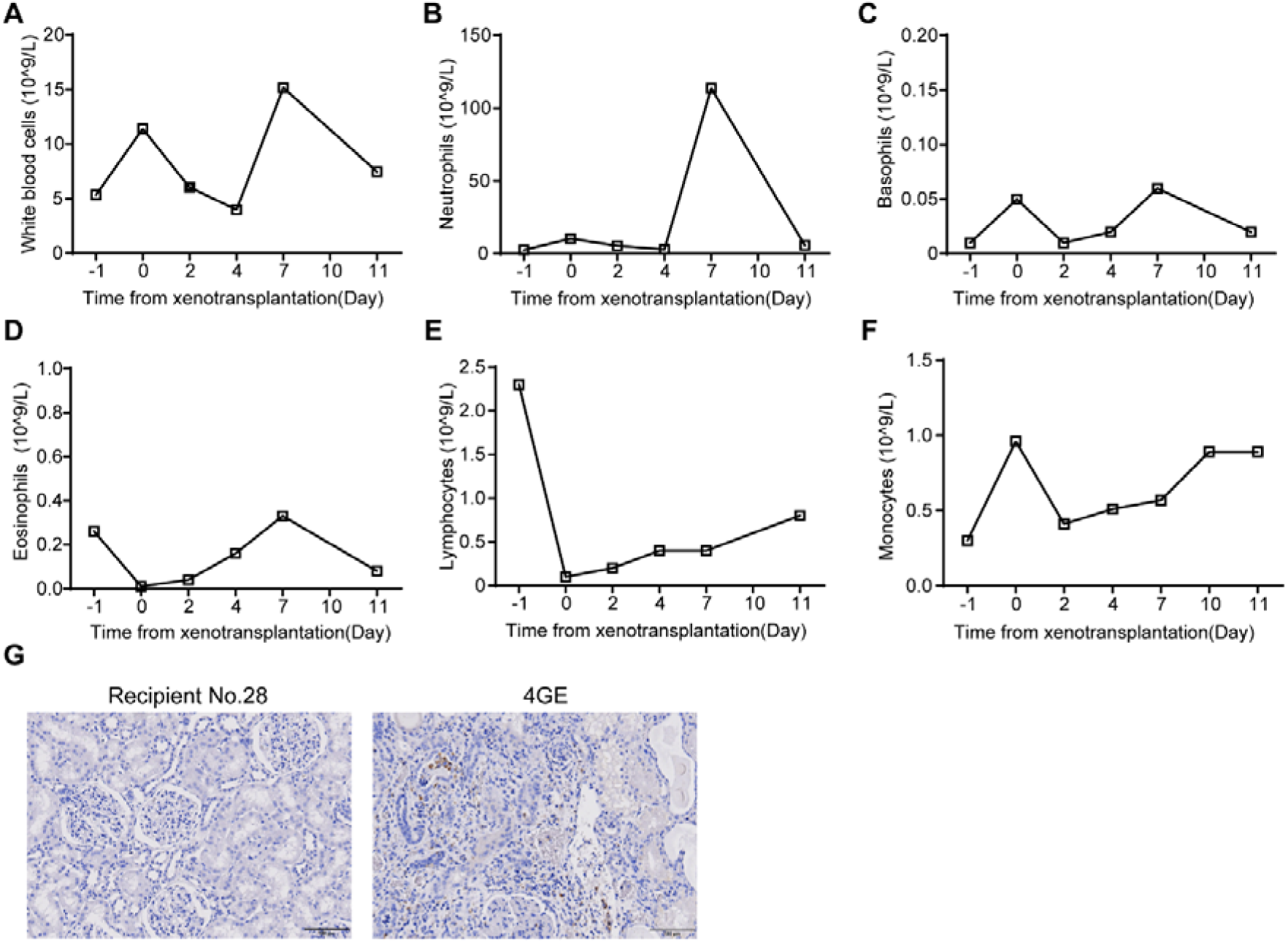
Cellular immunity of recipient monkey. A-F. The numbers of white blood cells (A), neutrophils (B), basophils (C), eosinophils (D), lymphocytes (E) and monocytes (F) in the whole blood of recipient monkey. G. Macrophage infiltration in porcine kidney xenograft confirmed by immunohistochemistry, scale bar=100 μm.

In terms of coagulation, the number of platelets on the day of renal xenograft function loss was significantly reduced (Figure 8A), but there were no significant changes in thrombin time (TT), prothrombin time (PT), activated partial thromboplastin time (APTT) or fibrinogen (FIB) in the recipient monkey (Figure 8B-E), indicating that overexpression of hCD39 and hTBM improved coagulopathy in the recipient.

**Figure 8.**
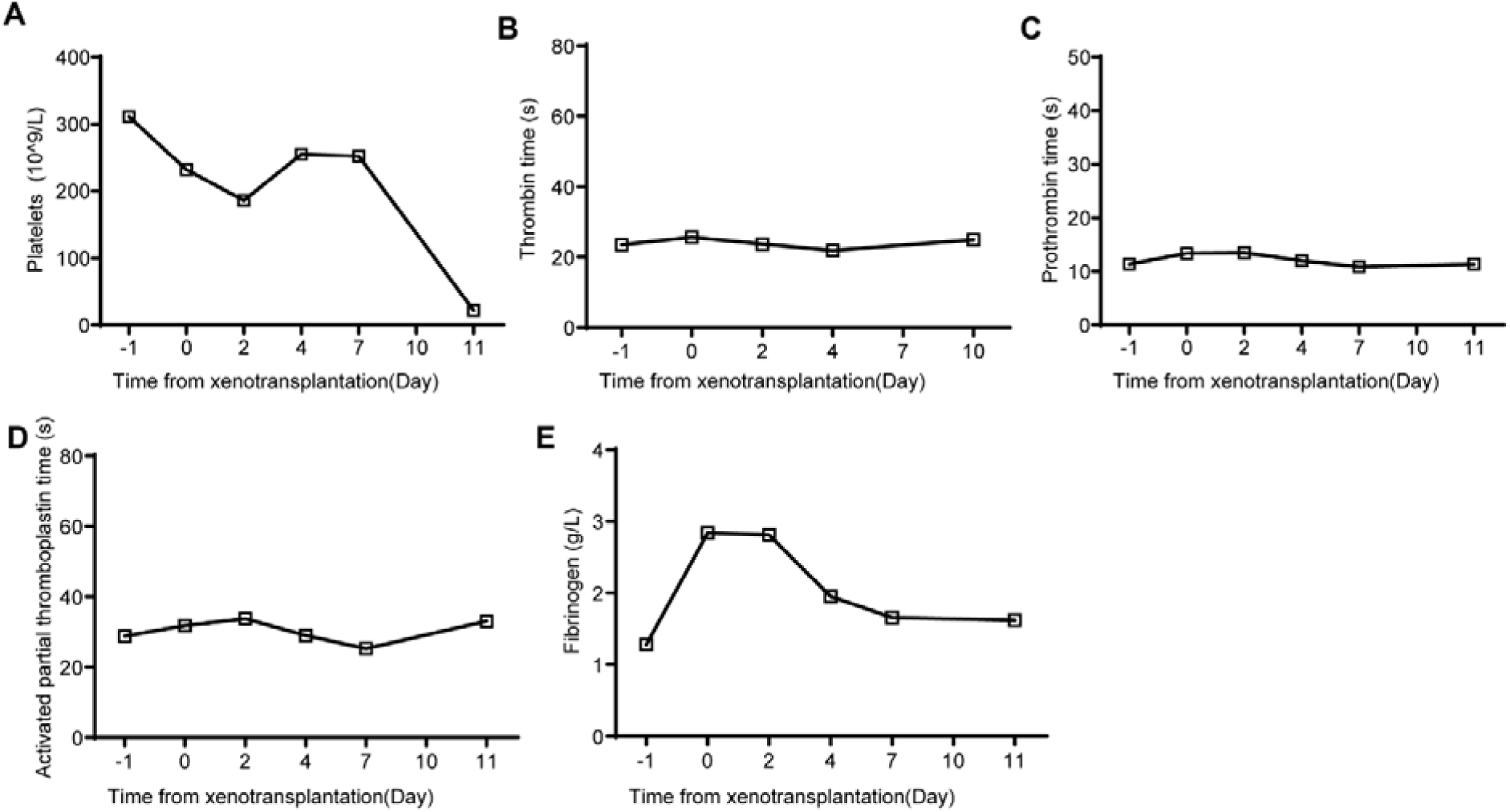
Coagulation function of recipient monkey. A. The numbers of platelets in the whole blood of recipient monkey. B-E. The coagulation indexes thrombin time (B), prothrombin time (C), activated partial thromboplastin time (D) and fibrinogen (E) in recipient monkey.

Physiological analysis revealed that the content of red blood cells (RBCs), hematocrit (HCT) and hemoglobin (HGB) of the recipient monkey were decreased after kidney transplantation. However, no significant changes in serum sodium, chloride, calcium, phosphorus, and potassium electrolytes were observed, while magnesium levels were significantly elevated on the day of functional loss of the kidney xenograft (Figure 9).

**Figure 9.**
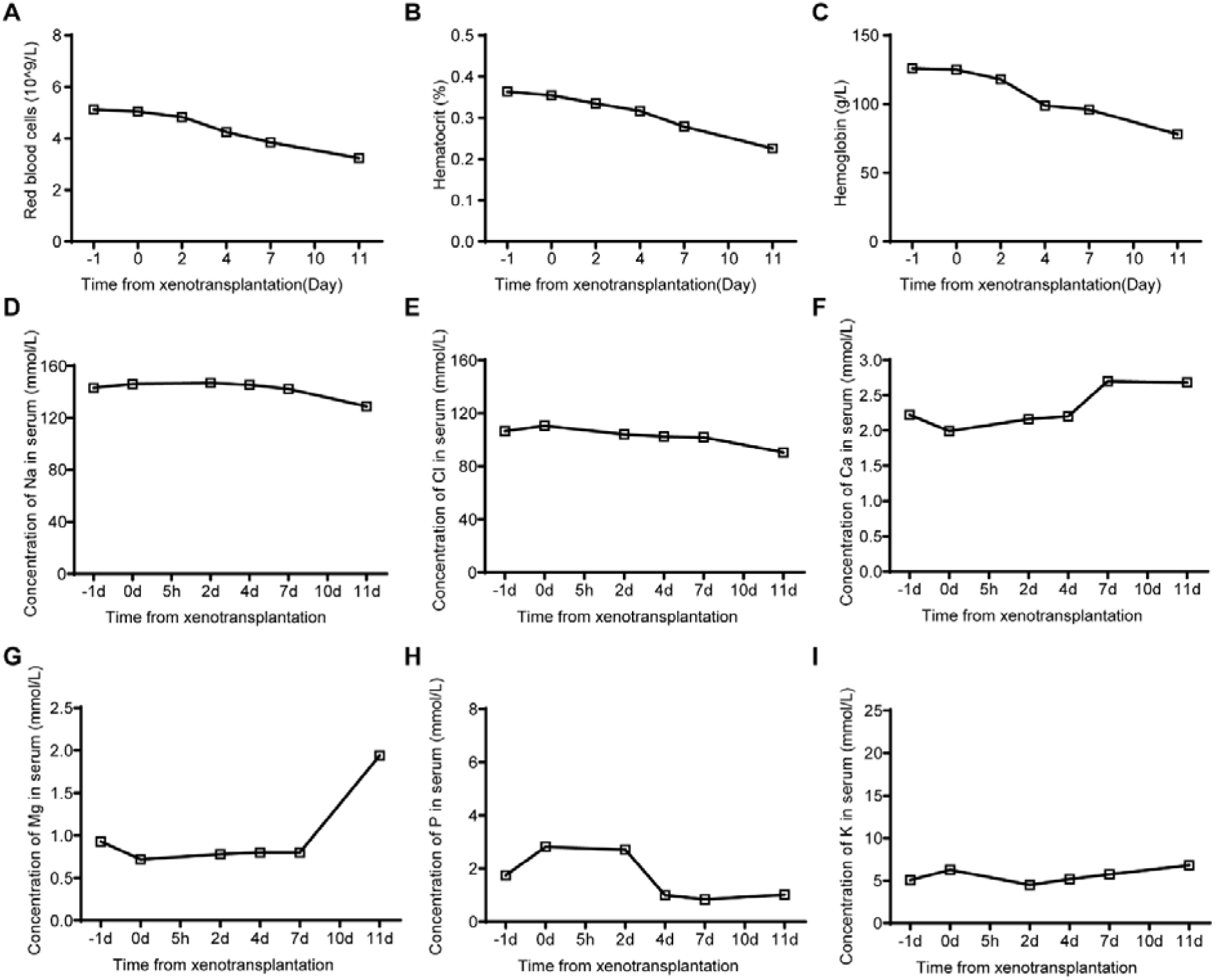
Erythrocyte indexes and electrolytes in recipient monkey. A. The numbers of red blood cells. B-C. Hematocrit (B) and hemoglobin concentration (C). D-I. The concentration of sodium (Na) (D), chlorine (Cl) (E), calcium (Ca) (F), magnesium (Mg) (G), phosphorus (P) (H) and potassium (K) (I) in the serum of recipient monkey.

In renal function tests, the serum creatinine, urea and cystatin C levels, urine creatinine levels, the ratio of urinary albumin and creatinine and urine microalbumin levels remained relatively stable in the recipient monkey within one week after xenotransplantation. However, on the day of renal graft function loss, these indicators increased significantly (Figure 10).

**Figure 10.**
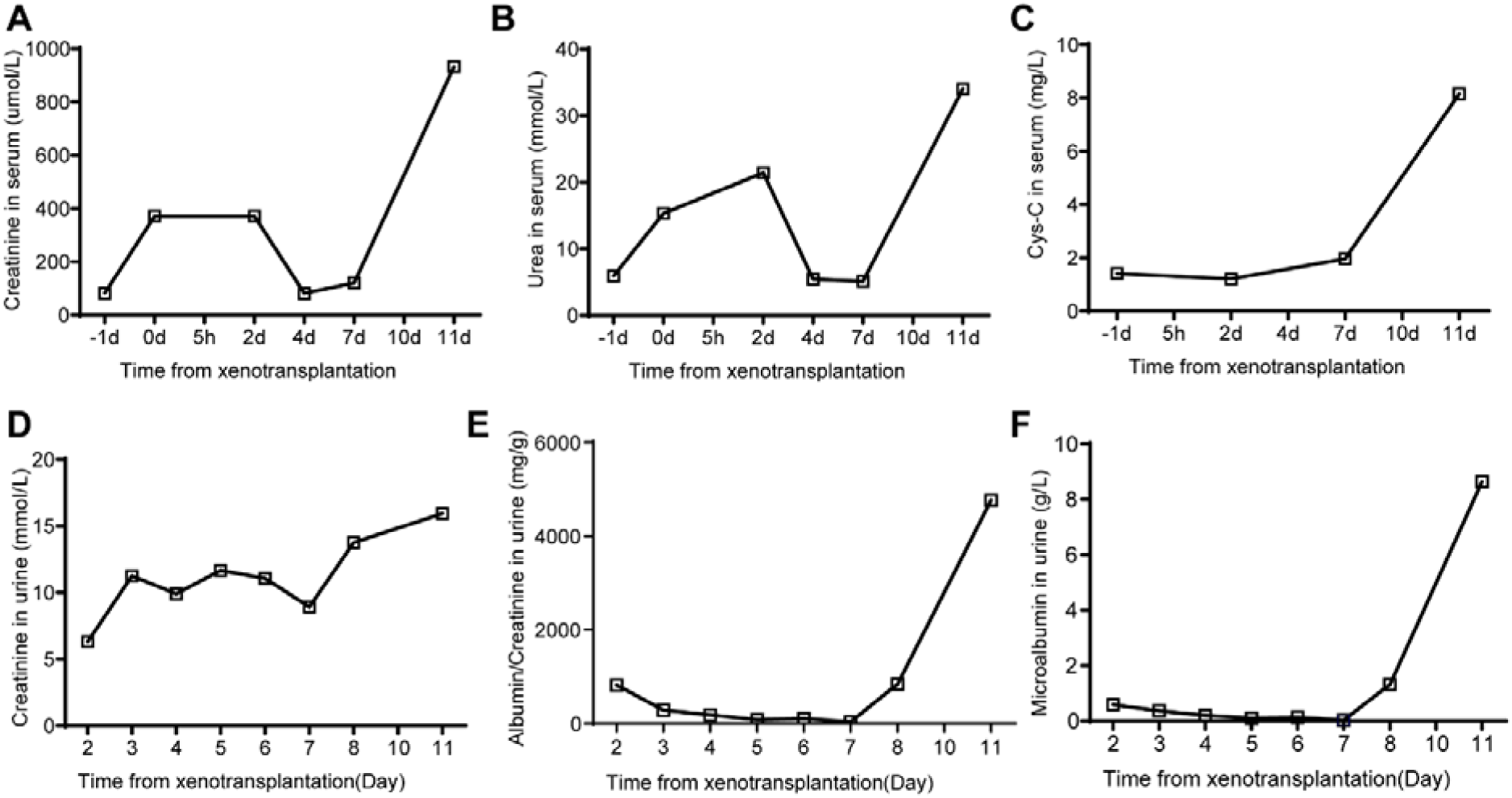
The renal function indexes of recipient monkey. A-C. The levels of creatinine (A), urea (B) and cystatin C (C) in the serum of recipient monkey. D-F. The levels of creatinine (D), ratio of ailbumin and creatinine (E) and microalbumin (F) in the urine of recipient monkey.

Liver function tests indicated that the total protein (TP), albumin (ALB), total bilirubin (TBIL), direct bilirubin (DBIL), alkaline phosphatase (ALP), aspartate aminotransferase (AST), adenosine deaminase (ADA) and alanine aminotransferase (ALT) levels also remained relatively stable within one week after transplantation. Two days after transplantation, total bile acid (TBA) levels were abnormally elevated in the serum of the recipient monkey. On the day of kidney graft function loss, TBIL and DBIL values in the recipient monkey were significantly elevated (Figure 11). These results indicate that our 4-GEC pig kidneys did not undergo HAR and functionally supported the life of the recipient monkey.

**Figure 11.**
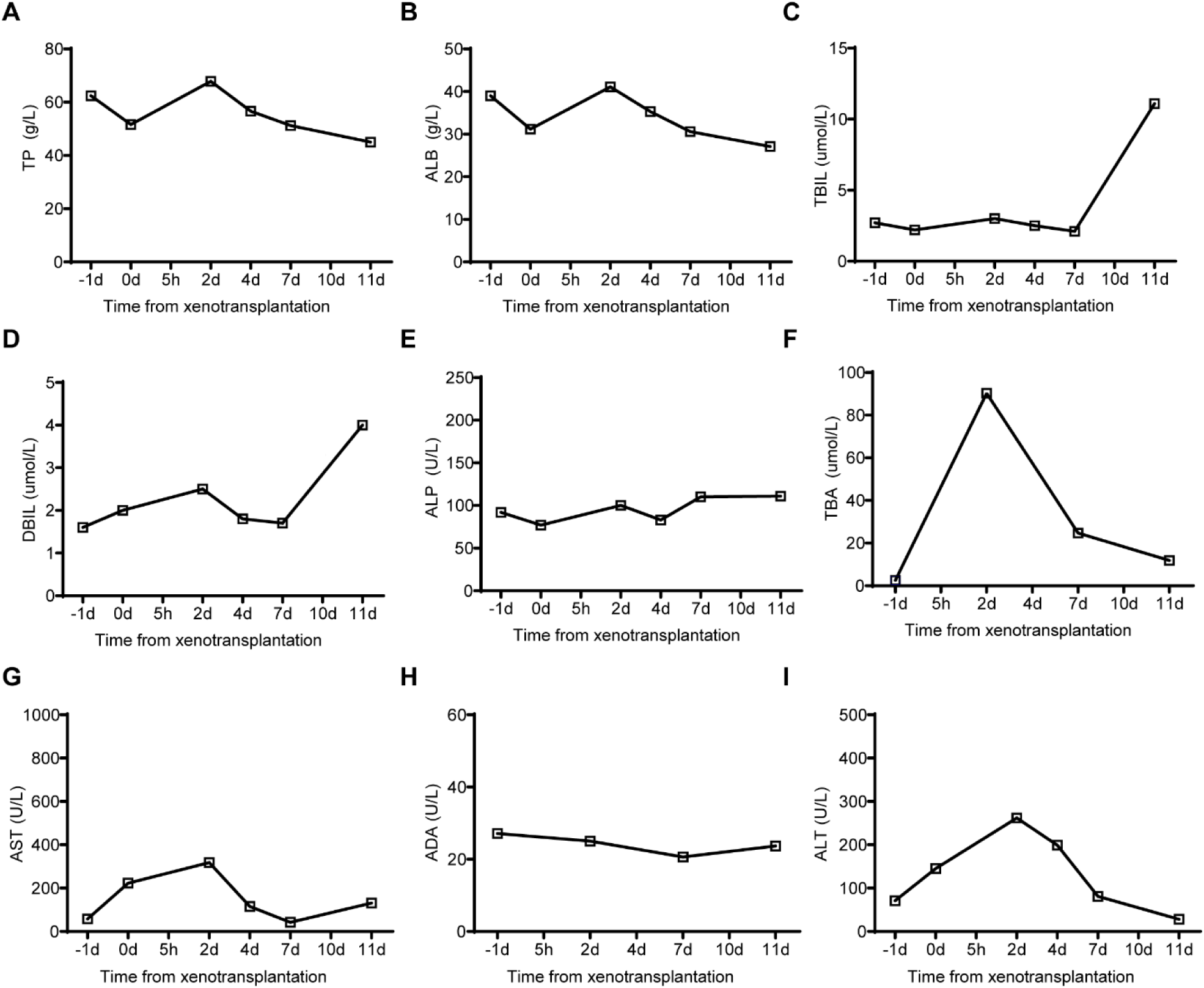
The liver function indexes of recipient monkey. A-I. The levels of total protein (TP) (A), ailbumin (ALB) (B), total bilirubin (TBIL) (C), direct bilirubin (DBIL) (D), alkaline phosphatase (ALP) (E), total bile acid (TBA) (F), aspartate transaminase (AST) (G), adenosine deaminase (ADA) (H) and alanine transaminase (ALT) (I).

## DISCUSSION

GEC pigs with GTKO [17], GTKO/hCD55 [18] and GTKO/hCD55/hCD39 [19] were produced, but the generation of GTKO/hCD55/hCD39/hTBM in the 4-GEC pigs has not yet been reported. In the production of the 4-GEC pigs, we simultaneously transfected 5 plasmids (Cas9 plasmid, 2 sgRNA plasmids, hTBM/hCD39 overexpression plasmids and transposon plasmids) into hCD55 transgenic *Diannan* miniature pig fetal fibroblasts. A total of 107 cell colonies were obtained by antibiotic screening, but most of them were not single-cell derived colonies. This is possibly due to the difficulty for a single cell to grow into a population after drug selection, and hence, they were prone to apoptosis. However, most of the cloning sites carried the hTBM/hCD39 transgene, indicating the high efficiency of the PiggyBac transposon system (Ding et al., 2005). To ensure that the GEC pigs have consistent genotypes and high transgene expression levels, we first obtained 7 cloned fetuses and conducted genotype identification and expression analysis of the fetuses. Among them, the F02 fetus had consistently high expression of hTBM, hCD39, and hCD55, and the F06 fetus had high expression of hTBM and hCD39. Therefore, we selected these two fetuses for recloning and obtained 58 live piglets. In addition, the expression of transgenes is usually inconsistent among different individuals even if these 4-GEC pigs are derived from the same FFCs [20]. Thus, we performed the third round of cloning by using individuals with high expression of each transgene and a normal phenotype.

The mRNA expression level of the GGTA1 gene in the 4-GEC pigs was significantly increased after GTKO (Figure 3B). This is consistent with our previous study and could be due to the regulatory mechanism of genetic compensation [21]. Nevertheless, flow cytometry and immunofluorescence analyses showed that the 4-GEC pigs did not express αGal (Figure 3C and D), suggesting that the GGTA1 gene was inactivated. In partial 4-GEC cells, the expression of hCD55 was undetected (Figure 3C). Therefore, we speculated that the partial region of the human CD55 sequence may be methylated due to reprogramming during the development of cloned fetuses, resulting in transgene silencing in some tissue cells [22].

In pig-to-NHP xenotransplantation studies, screening primates with low antibody titers as recipients is critical for xenograft survival [8, 23]. In this study, all of our GEC pigs had lower monkey IgG and IgM binding levels and higher CDC resistance than wild-type pigs (Figure 5A-C). We also used ATG and anti-CD20 mAb as induction therapy and an anti-CD154 mAb-based immunosuppressive regimen for maintenance therapy. However, the recipient monkey only survived for 11 days. Nevertheless, we found that most of the physiological indicators, immune levels, and kidney and liver functions were maintained at normal levels, and no HAR was observed during the postoperative monitoring of the recipient monkey.

The long-term survival of GE pig kidneys is mostly accompanied by the occurrence of thrombotic microangiopathy. In this study, we monitored coagulation in monkey and found that the recipient monkey did not develop coagulation disorders during their survival, further providing evidence that hCD39 and hTBM can effectively regulate coagulopathy. However, we also found that the recipient monkey rapidly developed thrombocytopenia before death, along with a small amount of complement C3c deposition in the kidney graft, which may be due to acute humoral immune rejection mediated by non-Gal antibodies.

### Conclusions

In summary, we successfully generated GTKO/hCD55/hTBM/hCD39 4-GEC xenotransplantation donor pigs for the first time and confirmed the anti-immune rejection and coagulation regulation effect of the 4-GEC pigs in the preclinical exploration of kidney xenotransplantation from pigs to NHPs.

## List of Abbreviations

ADA: Adenosine deaminase
ALB: Albumin
ALP: Alkaline phosphate
ALT: Alanine aminotransferase (ALT)
AST: Aspartate aminotransferase
ATG: Antithymocyte globulin
cDNA: Complementary DNA
COCs: Cumulus-oocyte complexes
DAMPs: Damage-associated molecular patterns
DBIL: Direct bilirubin
ddPCR: Droplet digital PCR
GE: Gene-edited
GEC: Gene-edited cloned
GGTA1: α1,3-galactosidyl transferase
NHPs: Non-human primates
NETs: Neutrophil extracellular traps
PBMCs: Peripheral blood mononuclear cells
PFFs: Pig fetal fibroblasts
RT: Room temperature
SCNT: Somatic cell nuclear transfer
SD: Standard deviation of mean
SNPs: Single nucleotide polymorphism
TBA: Total bile acid
TBIL: Total bilirubin
TMB: Thrombomodulin
TP: Total protein
4GE: GTKO/hCD55/hTBM/hCD39 gene-edited

## Declarations

### Consent for publication

Not applicable.

### Availability of data and materials

The dataset supporting the conclusions of this article is included within the article and its additional file.

## AUTHORS’ contributions

HJW and HYZ designed the research. CY, YW, XL, JW, XH and HZ performed the molecular experiments and analyzed data. DJ, ZX transfected the cell and performed a cell culture. JG, TW, YQ, HZ, and XZ carried out SCNT and embryo transfer. GC, CY, TW, and HZ performed pig-to-monkey kidney xenotransplantation. KX wrote the manuscript. HJW, HYZ and MAJ revised the manuscript and validated the data. All authors contributed to the article and approved the submitted version.

## Acknowledgments

We thank the Ministry of Science and Technology of the People’s Republic of China (Grant No. 2019YFA0110700) and the Department of Science and Technology of Yunnan Province (Grant No. 202102AA310047) for their financial support. We extend our thanks to Hui Yang (Shanghai Institutes for Biological Sciences) for gifting us PiggyBac vector and to Dr. Wang Gang (Department of Pharmacology Innovative Institute of Basic Medical Sciences of Zhejiang University) for providing anti-CD154 monoclonal antibody [5C8].

---

**Figure S1.**
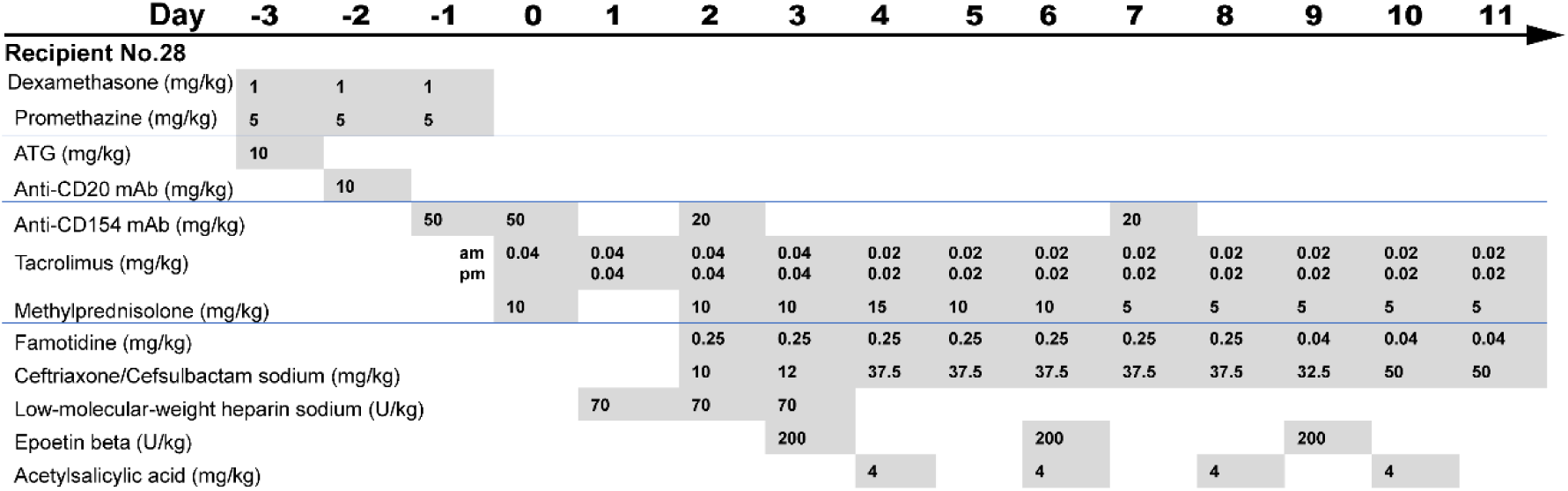
Immunosuppressive regimen.

**Table S1.**
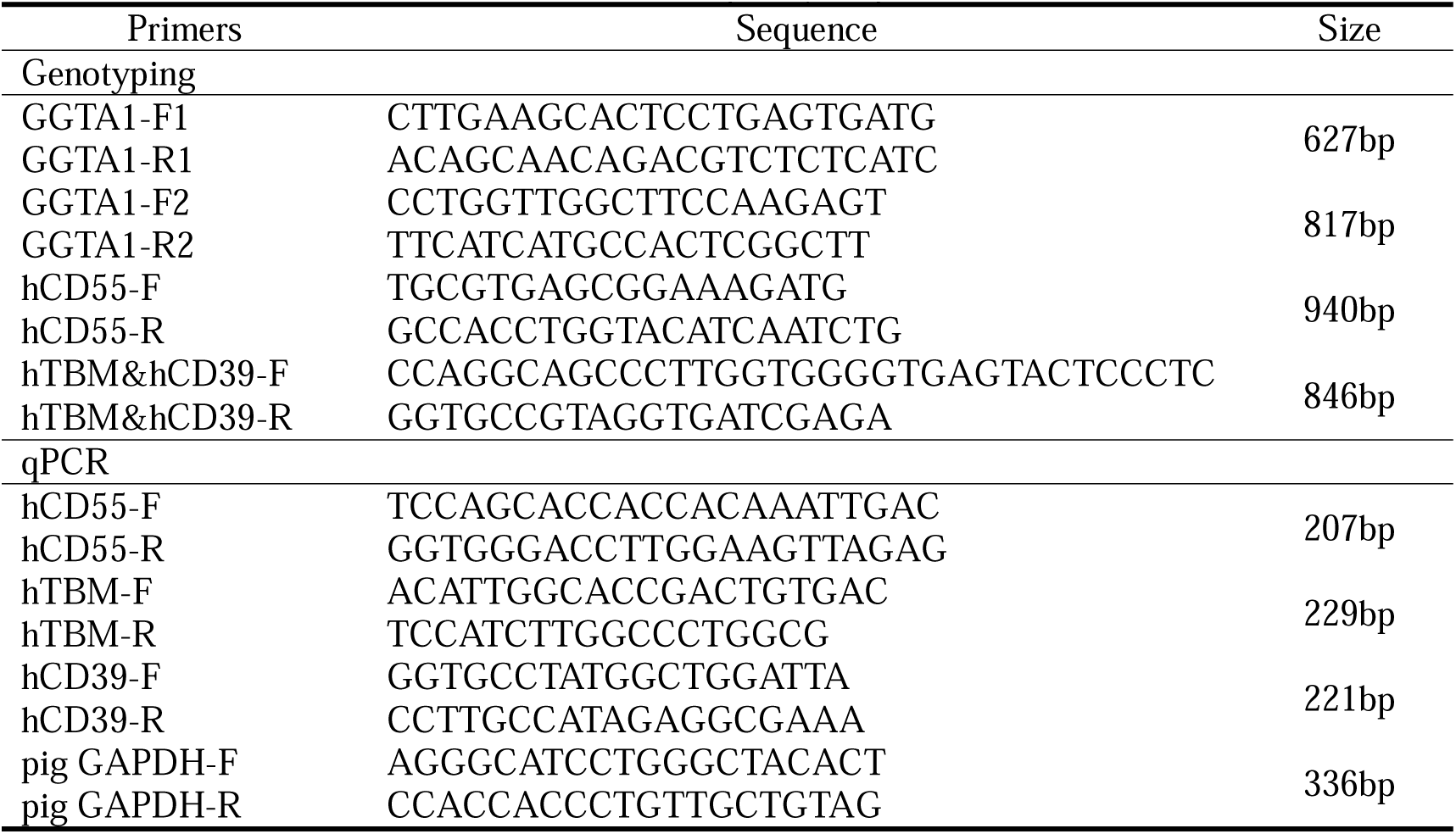
Primers for genotyping, qPCR and ddPCR.

**Table S2.**
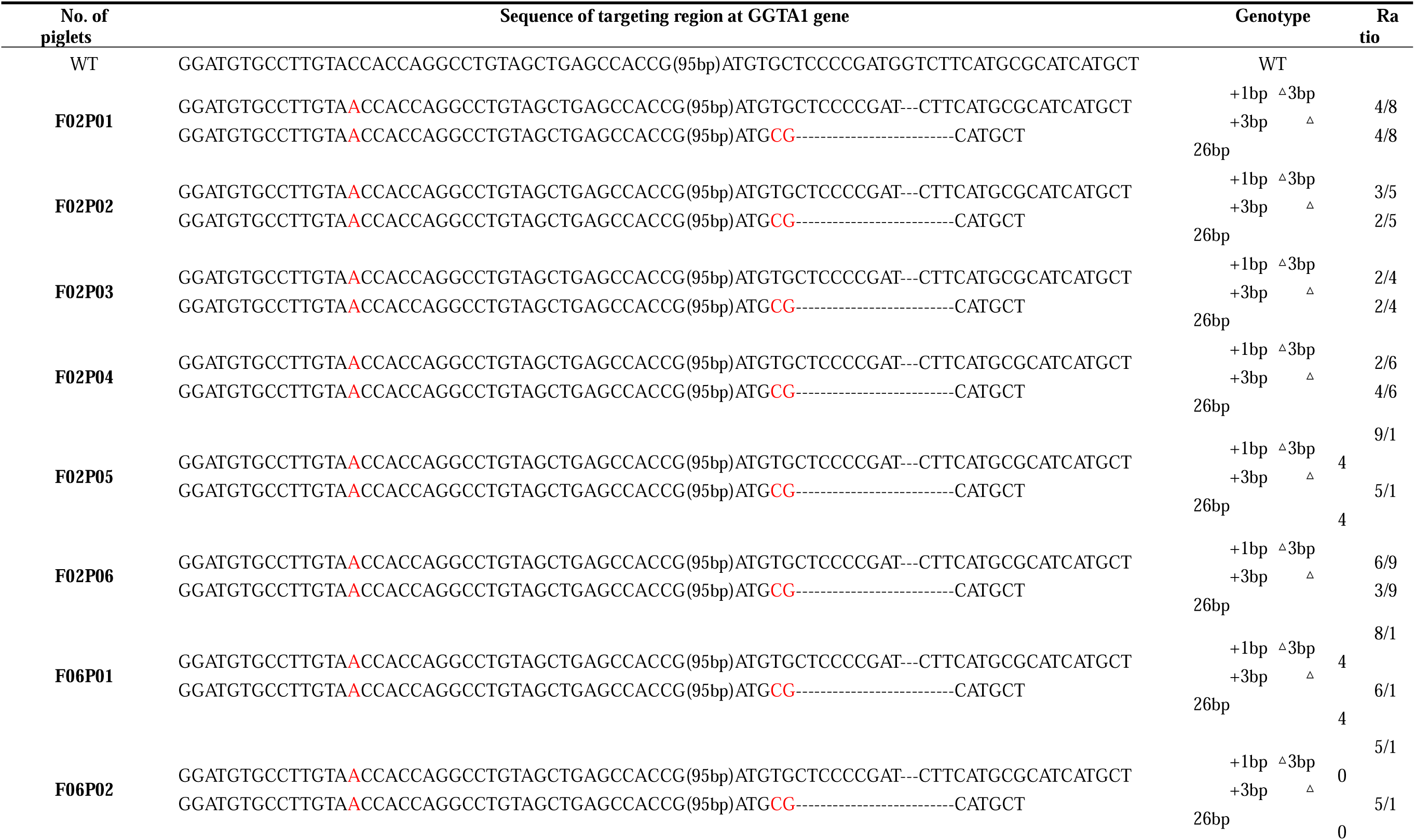
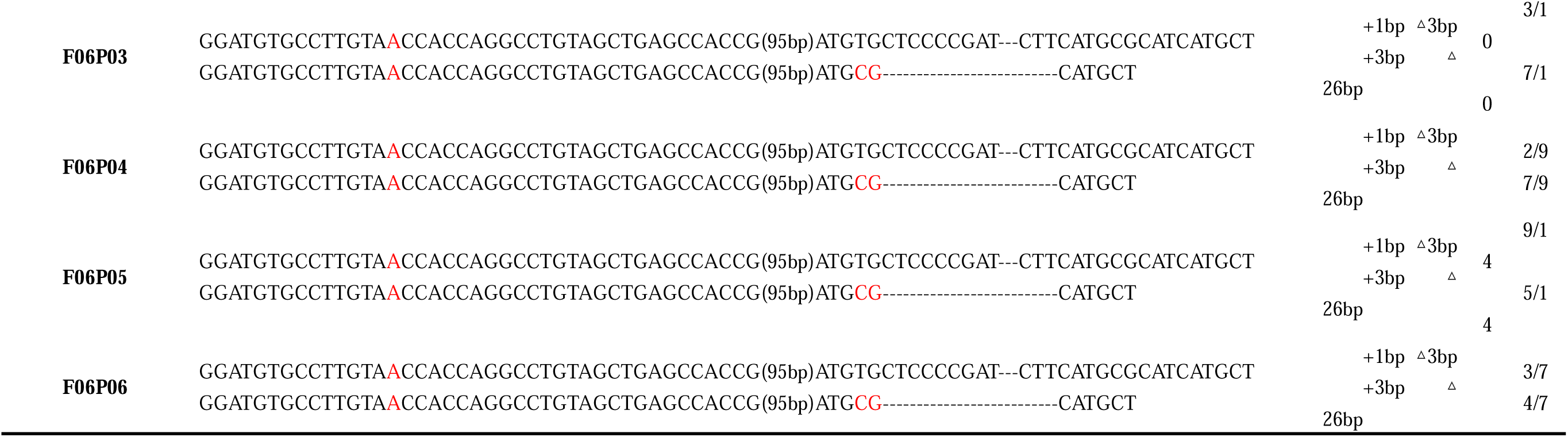
Genotype of targeting region at GGTA1 gene in cloned piglets derived from F02 and F06 fetuses.

